# An *in vitro* - agent-based modelling approach to optimisation of culture medium for generating muscle cells

**DOI:** 10.1101/2021.09.28.461963

**Authors:** David Hardman, Katharina Hennig, Edgar R Gomes, William Roman, Miguel O Bernabeu

**Affiliations:** Centre for Medical Informatics, Usher Institute, The University of Edinburgh, Edinburgh EH16 4UX, United Kingdom; Instituto de Medicina Molecular, Faculdade de Medicina, Universidade de Lisboa, Avenida Professor Egas Moniz, 1649-028, Lisboa, Portugal; The Bayes Centre, The University of Edinburgh, Edinburgh EH8 9BT, United Kingdom

## Abstract

Methodologies for culturing muscle tissue are currently lacking in terms of quality and quantity of mature cells produced. We analyse images from *in vitro* experiments to quantify the effects of culture media composition on mouse-derived myoblast behaviour and myotube quality. Computational modelling was used to predict an optimum media composition for culturing. Metrics of early indicators of cell quality were defined. Images of muscle cell differentiation reveal that altering culture media significantly affects quality indicators and myoblast migratory behaviours. To predict cell quality from early-stage myoblast behaviours, metrics drawn from experimental images or inferred by Approximate Bayesian Computation were applied as inputs to an agent-based model (ABM) of differentiation with quality indicator metrics as outputs. We describe cell behaviours as a set of functions of media composition to predict cell quality using the ABM. Our results suggest that culturing muscle cells in neural cell differentiation medium reduces cell-cell fusion but does not diminish cell quality and that increasing serum concentration increases myoblast fusion implying a trade-off between the quantity and quality of cells produced when choosing a culture medium. Our model provided a good prediction of experimental results for media with 5% serum provided the myoblast proliferation rate was known.

**Author summary:** Functional skeletal muscle tissue can be grown in the lab but is most useful if the constituent muscle cells behave as they would *in vivo*. Optimising the conditions to promote precursor muscle cell fusion and growth is therefore vital. With many different factors influencing cell growth finding optimal conditions through rounds of experimentation alone is difficult especially as we strive to complexify our cultures with multiple cell types. We created metrics quantifying mature muscle cell quality at an early stage of development and applied them to experiments with a variety of culture media compositions. Changing the concentration of serum and proportion of neuron differentiation medium produced differences in the behaviour of fusing cells and in the quality and quantity of mature cells. From these results we created phenomenological models describing the behaviour of fusing cells for any combination of serum and neuron medium concentration. We integrated these models into a multiscale agent-based computational model of cell fusion to predict cell quality and quantity through virtual experiments in an emergent fashion. Our model suggests that choosing culture media composition will involve fundamental compromises between cell quantity and quality and that cells which initially fuse quickly may produce less final yield.

## Introduction

A major challenge in the engineering of functional skeletal muscle tissue is the inability to reproduce the complex *in vivo* microenvironment of muscle tissue *in vitro* [1]. To form contractile myotubes, distinct cell sources are required. Currently, the most relevant *in vitro* myogenesis model employs primary myoblasts [2] and a tailored culture medium to successfully differentiate myoblasts into multinucleated myotubes.

Successful *in vitro* myogenesis is dependent upon cell culture medium composition and requires it to be fine-tuned to produce healthy and mature myotubes. Since *in vivo* myogenesis relies on several cell types, it is also important to ensure that the designed medium is compatible with these other cell types for muscle co-cultures.

Screening the media conditions that facilitate optimal muscle cell formation is necessary to maximise the quality, yield and reproducibility while reducing costs and experimental time. Relying on experimental trial and error to generate more sophisticated cell-based *in vitro* systems is impracticable and new strategies are necessary for culture medium optimisation [3].

*In vitro* experiments are invaluable tools for exploring cell culture environments [4] but they remain expensive, time consuming and provide sparse data points for analysis. Numerical models of cell and tissue behaviours [5] present fast, low-cost methods for simulating *in vitro* experiments but require thorough calibration against experimental results to establish confidence in predictions.

Given the lengthy duration of time required to produce mature myotubes *in vitro*, the ability to predict outcomes of cell experiments from early-stage indicators of cell behaviour would save experimental time and costs. Finding a single early-stage behaviour which is predictive of mature cell quality would allow direct inference of cell outcomes but if one is not apparent, a method for modelling the interactions between these early-stage behaviours would provide predictions of cell outcomes.

Agent based models (ABM) apply simple sets of behavioural rules to autonomous agents (such as cells or cell nuclei) and simulate their interactions with each other and their environment in order to recapitulate higher level emerging behaviours. ABM have been used to compliment cell and tissue engineering as a method to gain insight into the underlying mechanisms [6], but to our knowledge have not directly been applied to the challenge of optimisation of cell culture environments.

Studies using ABM generally derive agent rules from literature [7]. Here, we present a workflow for a combined *in vitro-in silico* approach in which experimental data is used to define behavioural metrics and apply these as rules to govern an ABM which can simulate cell quality from early stage behaviours and inform future *in vitro* experiments.

We define the cell culture variables to be optimised then image a limited set of *in vitro* trials over a range the chosen variables. We create metrics of appropriate indicators of cell quality from cell imaging, define metrics of early-stage agent behaviours from live imaging and select behaviours which show significant differences with changes in cell culture variables as potential predictive behaviours.

If no single early-stage behaviour can predict mature cell quality, an ABM is designed and calibrated to model the complex interactions of early-stage behaviours and produce metrics of cell quality indicators as an output. If the ABM can reproduce the measured cell quality indicators from *in vitro* trials, our goal is to create a predictive model of mature cell quality without the need to measure further behaviour metrics. This is achieved by interpolating the measured metrics of early-stage behaviours over all appropriate cell culture variable values and applying these as inputs to the ABM. ABM predictions are then validated by comparison of quality metrics with a further *in vitro* trial. Validation will determine whether the *in vitro - in silico* workflow described can predict cell outcomes based upon interpolated early-stage behaviour metrics alone or if further behavioural information is required.

We selected serum concentration as the first media variable since primary myoblast fusion into multinucleated myotubes occurs upon serum reduction [8] but the cellular effects of varying serum at these low concentrations remain ill-defined. We chose the proportion of neuronal media as our second component since co-culturing muscle cells with neurons increases muscle cell maturation through neuronal derived factors [9] and provides the possibility to study neuromuscular junction (NMJ) formation.

Healthy muscle cells are long multinucleated cells with evenly distributed nuclei throughout the periphery of the fibre. Since uneven or centralised distribution of nuclei is often linked to a disease state or improper maturation [10], [11], nuclear position and distribution is a good marker of muscle cell maturity and health. We therefore chose nuclei positions of both unfused myoblasts and fused myotubes (myonuclei) as the agents in our ABM. Nuclei are also suited to the role of agent here as their motion can be tracked over time from images, enabling quantification of their behaviour. This enables us to calibrate a model linking nuclear behaviour with cell health.

Our workflow for predicting muscle cell quality applies metrics of early stage cell behaviours to a nuclei-centred ABM with two phases, the first modelling myoblast motion and fusion and the second the nuclei force balance within myotube cells with metrics of cell quality indicators as an output. Key metrics of cell behaviours were observed from *in vitro* imaging and inferred using an approximate Bayesian computation sequential Monte Carlo (ABC-SMC) method. By extrapolating cell behavioural metrics from discrete experiments, we describe cell behaviours as a function of differentiation media composition which, when used as an input for a calibrated ABM, can produce *in silico* experiments which predict the range of media conditions in which muscle cells can thrive and highlight the optimum conditions for culturing muscle cells.

## Results

### Changes in differentiation medium composition affect both early behaviour and quality indicators

Generating bottom-up cell-based systems relies on the timely development of cells. Cellular behaviours early in the culture will therefore govern the culture’s quality at later stages. As such, we used live and fixed imaging to identify a list of potential indicators divided into early stage behaviours and quality indicators (Table 1). To be of use in a predictive model, indicators must show a significant variation in relation to one or more of the cell culture variables being optimised.

**Table 1.** List of Metrics.

| Metric | Early stage behaviour (B) or quality indicator (Q) | Used in workflow |
| --- | --- | --- |
| Myoblast speed of motion | B | Used |
| Myoblast variation in angular velocity | B | Used |
| Myoblast rate of proliferation | B | Used |
| Myoblast angular velocity | B | Not used |
| Myoblast persistence of motion | B | Not used |
| Myoblast velocity | B | Not used |
| Myonuclei density | Q | Used |
| Myonuclei coefficient of variation | Q | Used |
| Mean distance between myonuclei | Q | Used* |
| Myotube width | Q | Not Used |
| Proportion myotubes with actin striations | Q | Not Used |

\*Applied as an accompanying indicator to coefficient of variation.

Culture medium is a central lever in influencing cell quality. We varied levels of serum and neuronal medium independently and observed significant changes in both early cell behaviour and quality indicators.

Potential indicators of mature cell quality were extracted from fixed, stained images(Fig 1A). The difference in myonuclei per mm^2^ between days 0 and 5 provides an indication of the total amount of cell fusion occurring during differentiation. One hallmark of mature myotubes is the presence of multiple myonuclei, located at the cell periphery, uniformly distributed throughout the fibre. Uniformity of myonuclei distribution was evaluated using the coefficient of variation, calculated by dividing the standard deviation of distances between nuclei in a cell by the mean distance between nuclei in the respective cell. A lower coefficient of variation indicates greater uniformity. Using spatial uniformity of myonuclei as an indicator of quality assumes that nuclei are spread sufficiently far apart. As the coefficient of variation does not provide information on distances between nuclei, the mean distance between myonuclei was also recorded.

**Fig 1.**
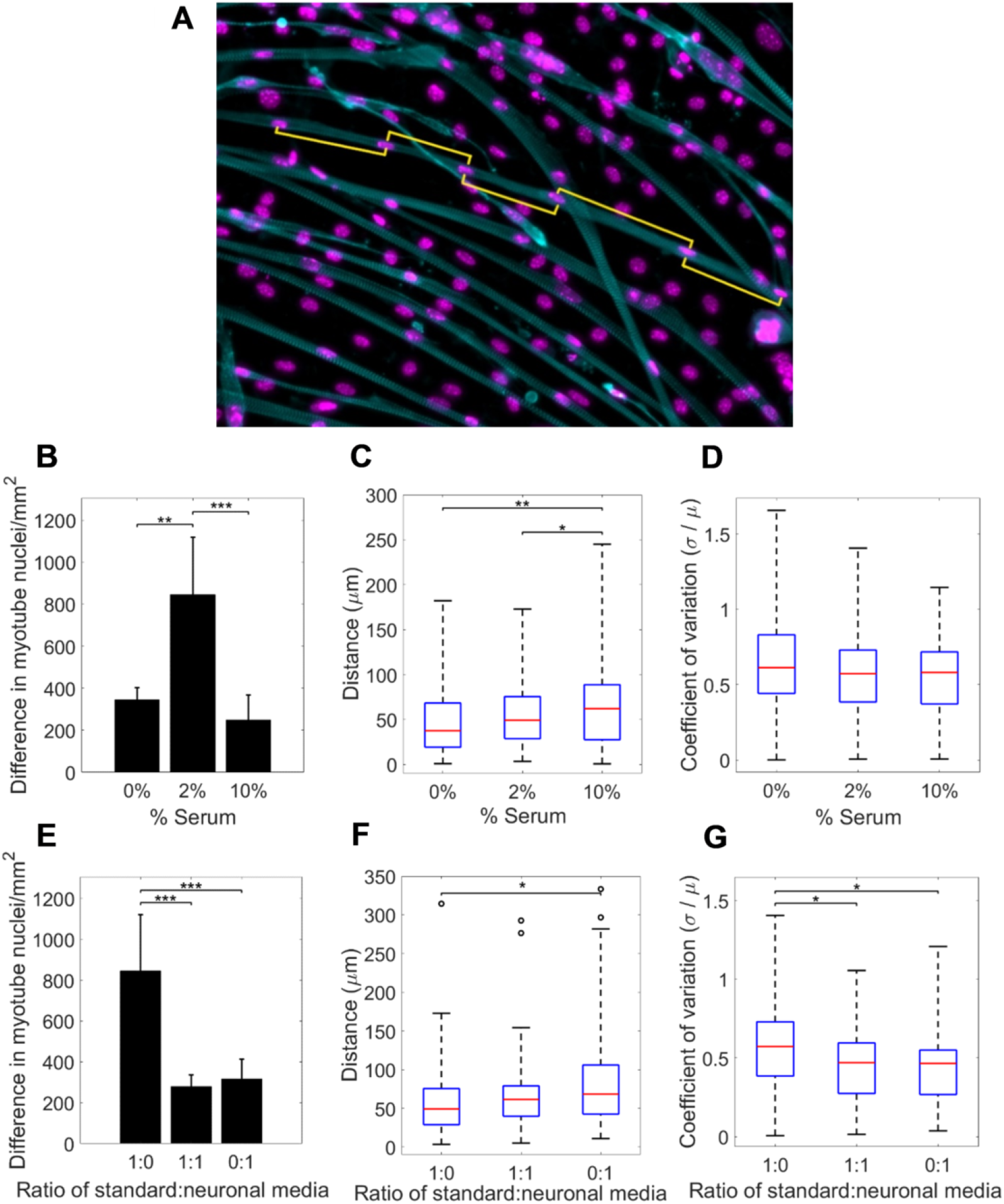
Myotube quality indicators at day 5 of differentiation. (A) A fixed image stained for cell nuclei (magenta) and actin cytoskeleton (cyan), yellow lines indicate measured distances between myonuclei. Plots (B-D) show differences in quality indicators with varying serum concentrations. (B) Difference in myonuclei per mm^2^ between days 0 and 5 of differentiation, (C) mean distance between myonuclei and (D) uniformity of distance between myonuclei expressed as ‘coefficient of variation’. Plots (E-G) show differences in quality indicators in varying ratios of neuronal medium. (E) Difference in myonuclei per mm^2^, (F) mean distance between myonuclei and (G) coefficient of variation.

Increasing the serum concentration (Fig 1B) from 0% to 2% led to a significant (464 more myonuclei/mm^2^) increase in the density of myonuclei for cells cultured in muscle cell differentiation media. A further increase to 10% serum concentration resulted in a reduction in myonuclei density (559 less myonuclei/mm^2^) to a level slightly lower than with no serum. This suggests an optimal serum concentration to promote muscle cell differentiation *in vitro*. Increasing the concentration of serum in differentiation medium resulted in a gradual increase in the mean distance between myonuclei (Fig 1C) but, while there is some evidence of a reduction in the range of coefficient of variation, there was no significant change in uniformity (Fig 1D) of myonuclei distribution.

A significant drop in the density of myonuclei (567 less myonuclei/mm^2^) was observed between experiments cultured in muscle cell differentiation media and in muscle and neuronal cell differentiation media mixed in a 1:1 ratio (Fig 1E). Density of myonuclei remained at a similarly low level in experiments with 100% neuronal medium, demonstrating the existence of an optimal base medium for myotube formation. Increasing the proportion of neuronal medium lead to an increase in mean distance between myonuclei (Fig 1F). Experiments with 100% neuronal medium and a 1:1 mixture of neuronal to muscle cell differentiation media exhibited a lower coefficient of variation, and therefore more uniform distribution, than muscle cell differentiation medium alone (Fig 1G).

Mean thickness of myotube cells (Supplementary Materials Fig S1) and proportion of sampled myotubes with full or partial actin striations (Supplementary Materials Fig S2) were also studied as potential quality indicators, but did not display significant differences between trials with different media compositions and so were discounted as appropriate quality indicators for our workflow.

To create metrics of early stage cell behaviours, myoblast cells were segmented and tracked from live imagestaken during days 0-1 after the application of differentiation media (Fig 2A). Significant and non-linear differences in metrics of myoblast migration speed, the distribution of cell angular velocities and cell proliferation rates were observed.

**Fig 2.**
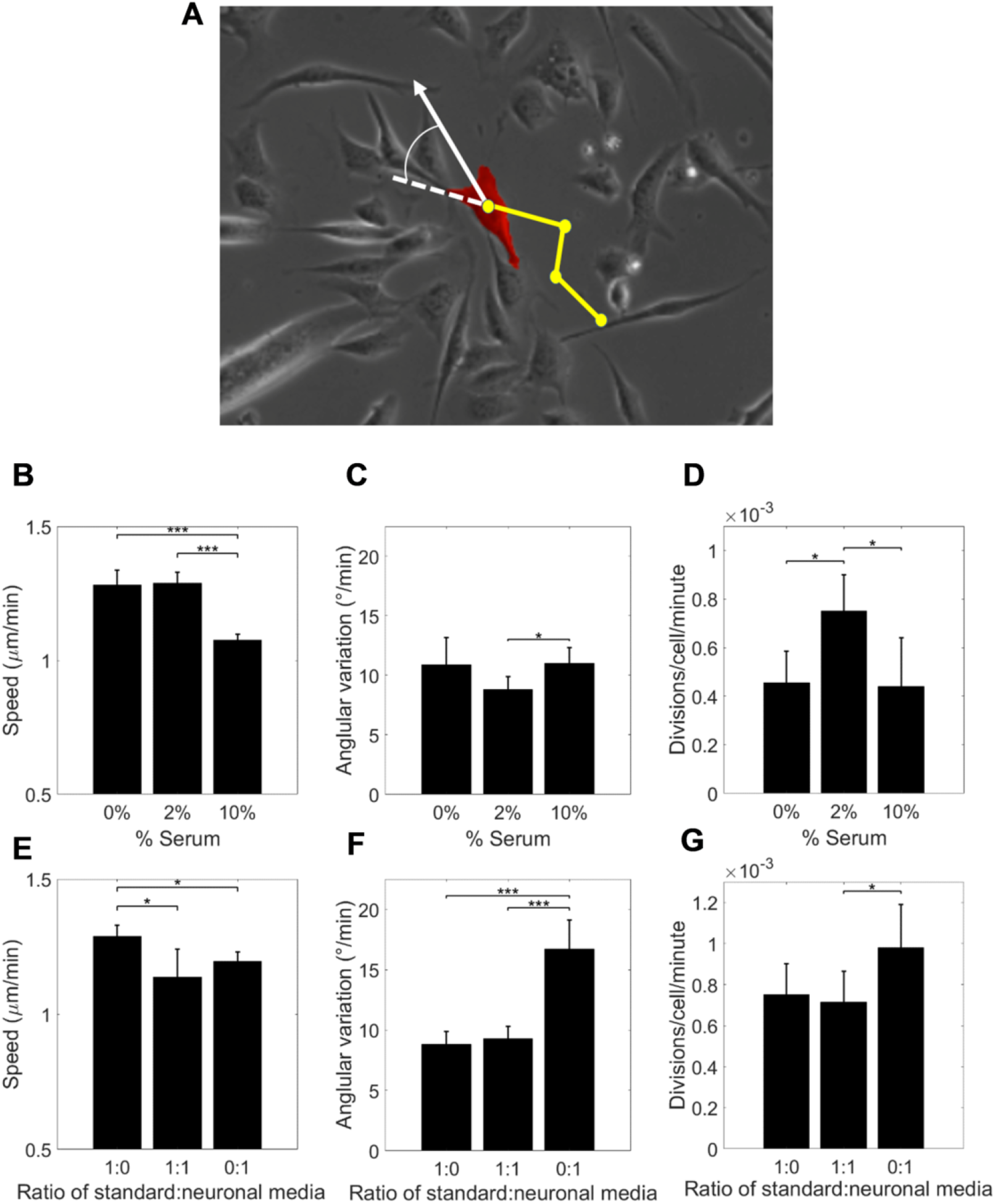
Metrics of myoblast behaviour from live images of cells. (A) Illustration of tracking a single myoblast (red) in a brightfield image. Yellow dots show position in previous frames. Plots (B-D) show differences in early-stage behaviour metrics with varying serum concentrations. (B) Myoblast speed, (C) standard deviation of angular velocity and (D) rate of myoblast division. Plots (E-G) show differences in behaviour metrics in varying ratios of neuronal medium and 2% serum concentration. (E) Myoblast speed, (F) standard deviation of angular velocity and (G) rate of myoblast division

For serum concentrations between 0-2%, myoblast cells moved at similar speeds(Fig 2B). Increasing serum concentration from 2% to 10% resulted in a significant reduction in cell speed. There was also evidence of a modest dip in the deviation of angular velocity as serum concentration is increased from 0-10% (Fig 2C). Proliferation rate of myoblasts increased significantly at a serum concentration from 2% (Fig 2D).

Concerning distinct muscle-neuron media mixtures, we observed a decrease in myoblast speed when muscle medium was replaced with neuronal differentiation medium or mixed to 1:1 ratios (Fig 2E). However, angular velocity (Fig 2F) and myoblast proliferation rate (Fig 2G) was only altered when muscle medium was fully replaced with neuronal medium. These data indicate that varying the media composition produces quantifiable changes in the behaviour of myoblast cells.

Mean direction, speed and persistence of myoblast motion were also extracted from tracking cells in live images. On average, cells in all images analysed displayed no global directional preference and so mean direction was discounted as an early stage behavioural indicator. While velocity and persistence of motion varied with media composition, we reasoned that these behaviours can be recapitulated using speed and angular velocity and so were not included to simplify the ABM inputs.

### A calibrated agent-based model predicts indicators of cell quality from *in-vitro* early-stage cell behaviours measurements

While our results show that both early behaviour and quality indicators vary with differentiation media composition, we found no direct relationship between any single early behaviour and the quality of mature myotubes. To study whether mature cell quality can be determined from the interaction between early-stage behaviours, we designed an ABM of myoblast fusion and myonuclei migration to simulate the complex interactions between early-stage behaviours and produce metrics of quality indicators as an output. The ABM is described in detail in Methods. Briefly, cell nuclei were used as the interacting agents in the model and a residence time mechanism was applied to simulate cell-cell fusion. Measured metrics of myoblast motion from live imaging were used to create distributions of behaviours to apply as inputs to the model. Fixed imaging was used to provide measurements of nuclei spatial distributions for initialising the model and for determining quality indicators for comparison. Calibration of the residence time mechanism and force-balance acting on fused nuclei requires the input of further metrics of cell behaviours which were not directly measurable from our imaging data (see Table 4 in Methods). The use of approximate Bayesian computation (ABC) allows us to infer distributions of these metrics by strategically running ABM simulations and comparing their outputs to measured quality indicator metrics. Two extra *in vitro* trials were conducted to provide conditions with further combinations of serum and neuronal differentiation medium. One trial with 5% serum and muscle and neuronal differentiation medium mixed in a ratio of 1:4 and the other with 10% serum and exclusively neuronal differentiation medium.

ABM simulations reproduced mean myonuclei densities to within 1 standard deviation of those observed experimentally for most conditions (Fig 3A). The exceptions of 2% serum and 1:1 muscle to neuronal cell differentiation media and 2% serum and 100% neuronal cell differentiation medium exhibit day 5 myonuclei densities significantly smaller than the initial number of myoblasts. The ABM assumes that myoblasts do not die during the experimental timeframe and all have the potential to fuse throughout the differentiation stage and so the total number of nuclei at day 0 is the minimum number of myonuclei obtainable from simulation.

**Fig 3.**
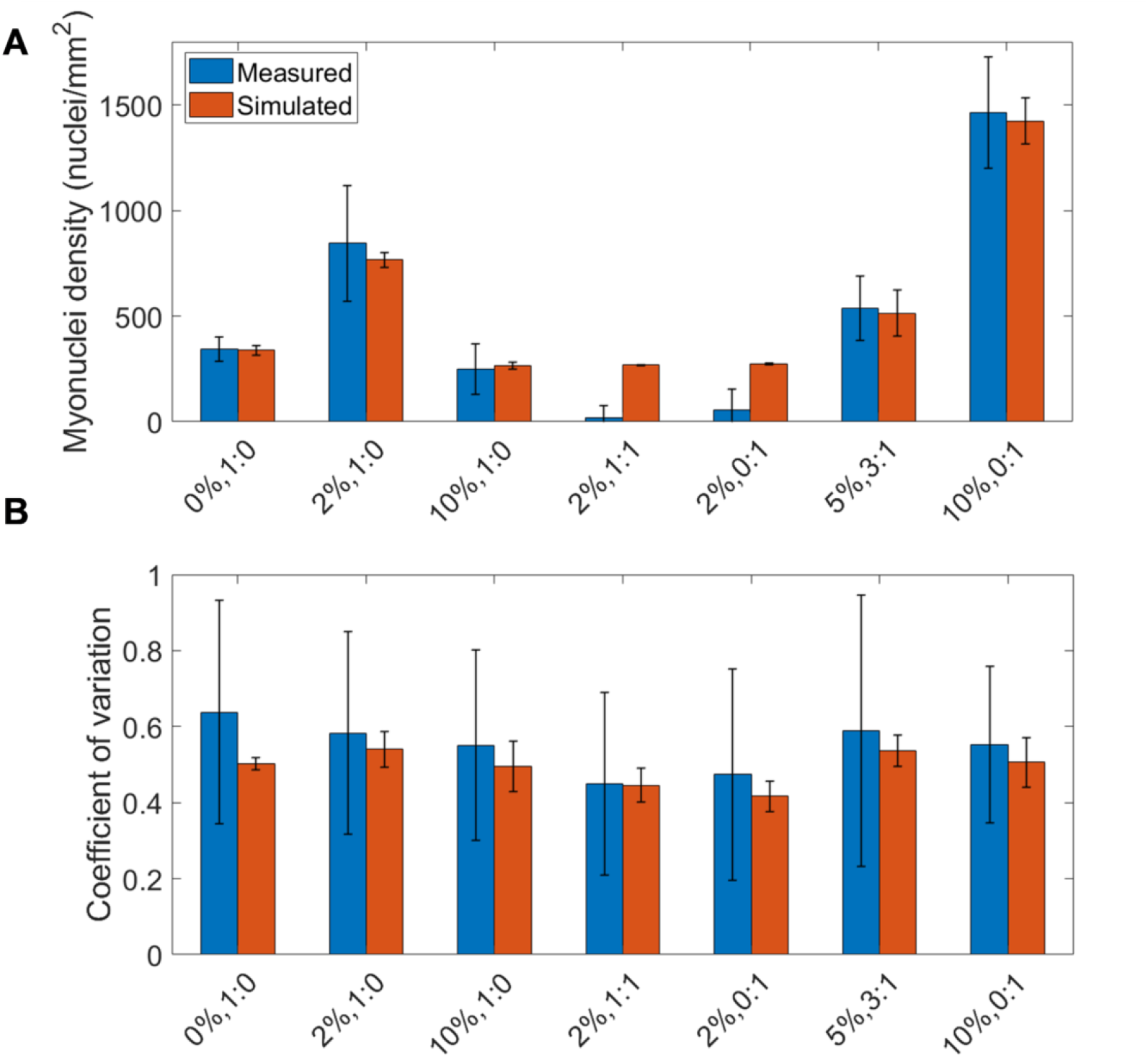
Comparison of measured and simulated quality indicators at day 5 of differentiation. Myonuclei quality metrics measured from imaging data from *in vitro* trials against outputs from the calibrated Agent-Based Model for varying serum concentrations and ratios of muscle to neuronal medium. (A) Myonuclei density. (B) Coefficient of variation, a metric of nuclei spatial uniformity.

To assess whether the ABM simulations successfully reproduce changes in quality indicators over time, the mean myonuclei densities in an *in vitro* 2% serum and no neuronal medium trial were measured on each day of differentiation and compared with simulations. Simulated myonuclei density (Fig S3) was found to be within 1 standard deviation with measured for each day.

Median values of coefficient of variation were reproduced by the ABM to within 1 standard deviation for all trials (Fig 3B and Supplementary Materials Table S1). The ABM results showed a smaller variation in myonuclei spatial uniformity than measurements from imaging data (Fig 3B and Supplementary materials Table S1). ABM simulations also reproduced mean distance between myonuclei per cell to within 1 standard deviation (Supplementary Materials Table S1)

These results imply that the *in vitro* / ABM workflow outlined here is sufficiently calibrated to reproduce average values of quality indicators in trials in which fusion occurs.

### Fitting surface models from discrete in-vitro experimental data enables prediction of myoblast and myotube behaviours

Our goal is to use the calibrated ABM to create a predictive model of muscle cell quality based on key early-stage cell behaviours without the requirement for further measurements of behaviour metrics. To achieve this, we first describe how these key behaviours change with respect to the composition of the differentiation media used. We assume that the effects of media composition on cell behaviour, while non-linear, are of a lower order than the complex effects of media composition upon cell quality. A second-degree polynomial surface model was applied to fit metrics of cell behaviours observed from live imaging (*S*_*mb*_, *ω*_*mb*_, and P) and inferred via the ABC-SMC method (*γ, t*_*MT fuse*_, *k*_*lat*_ and *k*_*nuc*_) to provide input cell behaviours for all concentrations of serum and neuronal medium. These predictions of cell behaviour are visualised in Fig 4A-F.

**Fig 4.**
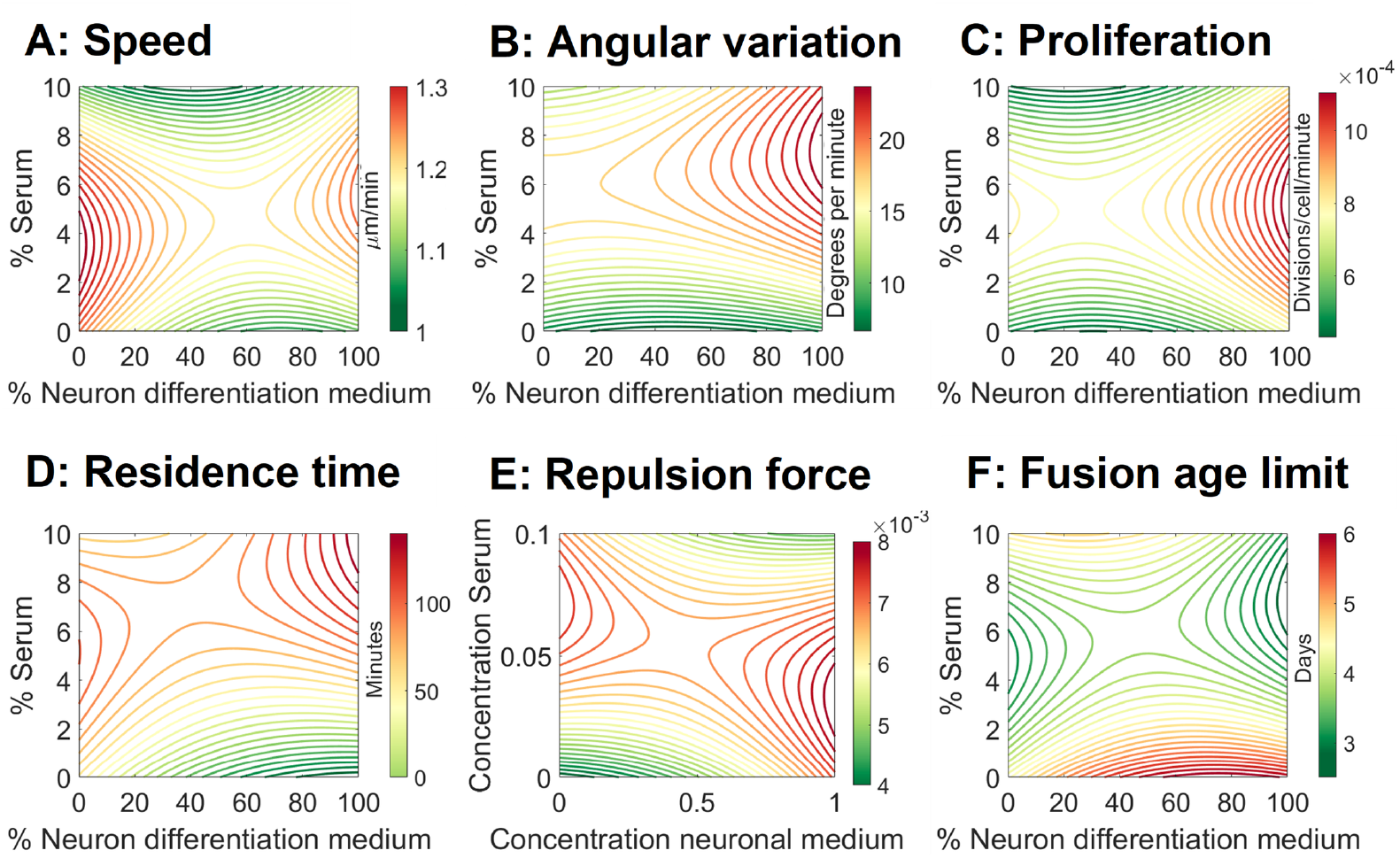
Fitted surface models for cell behaviours from *in vitro* imaging data. Second-order fit of observed (A-C) and inferred (D-F) behavioural parameters. (A) Speed of myoblast motion, (B) variation in myoblast angular velocity, (C) myoblast chance of proliferation, (D) residence time threshold, (E) nuclei repulsion force coefficient and (F) maximum age at which myotubes can fuse.

Maximum myoblast speed is projected to be found in media with 3.5% serum and no neuronal medium present (Fig 4A). An additional local maxima is shown to occur in media with 5.5% serum and 100% neuronal medium. Myoblast angular velocity (Fig 4B) shows a general increase in variation (and thus a decrease in cell persistence) with increased serum concentration, with a maximum of 23 degrees per minute occurring in media with 7.5% serum and 100% neuronal medium. A maximum chance of proliferation of 1.1×10^*−*3^ per cell per minute is expected for media with 5% serum and 100% neuronal medium (Fig 4C).

Increasing serum concentration was shown to correspond with an increase in the maximum residence time threshold (Fig 4D) suggesting that, at higher serum concentrations, myoblasts require a longer time in contact with myotubes before fusion can occur. The coefficient of myonuclei repulsion force (*k*_*nuc*_), (Fig 4E) showed a general trend for increasing with increase in serum concentrations, with a slight decline at high concentrations of serum (*>*~ 7%). For media with lower serum concentrations (*<* ~4%), *k*_*nuc*_ increases as the concentration of neuronal medium is increased.

The maximum age at which two or more myotubes may fuse together was projected to be between 3 to 4.5 days of cell differentiation for most media conditions (Fig 4F). This is in agreement with observations of actin striations occurring in cells after day 3. Myotube-myotube fusion was shown to end earlier in cells with increased levels of neuronal medium and continue for longer in cells with higher serum concentrations. Since simulations were ended at day 5, cells in a medium with no neuronal medium and 10% (or greater) serum concentration may still allow fusion at day 5 and beyond.

Fitting surfaces to these early stage behaviour metrics provides functions for estimating their values for any serum or neuronal media composition. These values can be applied as inputs to the ABM in order to predict myotube quality indicators throughout the parameter space.

### Difference in total myonuclei production is inversely related to early-stage nuclei fusion index

One advantage of ABM methods compared to regression models is the insight they give into the possible mechanisms which affect the outcomes of a model. From ABM simulations throughout the parameter space, it was noted that conditions generating small (300-400 nuclei/mm^2^) total increases in myonuclei by day 5 (Fig 5A) exhibit a negative exponential curve in the increase in myonuclei density over time and a steep initial decline in myoblast numbers while larger total increases in myonuclei (Fig 5B) are associated with a sigmoidal increase in myonuclei over time with nuclei initially increasing or remaining stable before decreasing. To account for these distinctive trends, we first compared the rate of myonuclei production between days 0-1 with day 5 difference in myonuclei for all media type experiments but found no significant correlation.

**Fig 5.**
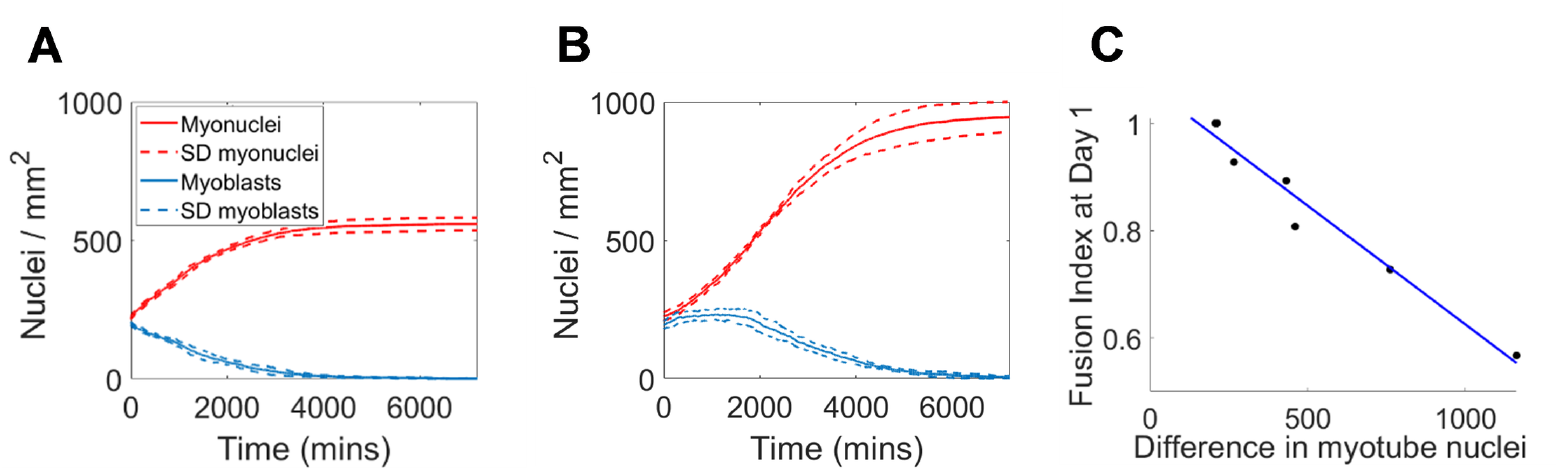
Relation between increase in myonuclei density and Fusion Index. Examples of Agent Based Model (ABM) generated changes in myonuclei (red) and myoblast (blue) nuclei density over time for conditions (A) 0% serum in neuronal medium and (B) 2% serum in neuronal medium (dashed lines represent standard deviation over 5 runs). (C) ABM generated differences in number of myonuclei between days 0 and 5 against Fusion Index (*R*^2^=0.96).

Further analysis indicates that these differences in total number of myonuclei (and therefore the number of fusion events) relate to the ratio of myonuclei to total nuclei, described by the nuclei fusion index [12], during the initial stages of differentiation. There is a strong negative linear relation (*R*^2^=0.96) between day 0 – day 5 difference in myonuclei density and the nuclei fusion index at day 1 (Fig 5C).

### ABM simulations predict a compromise between cell quantity and quality when selecting optimal culture media composition

Using the calibrated ABM to conduct a sweep of media compositions elucidates the variation in quality indicators and allows us to predict media compositions which will produce the optimum muscle cell quality.

Our model predicts that difference in myonuclei density at day 5, an indicator of total cell fusion, initially increases as serum concentration increases (Fig 6A). We also predict a general reduction in the difference in myonuclei density with increasing proportion of neuronal medium. These effects were shown to be non-linear, with maximum total fusion events predicted at serum concentrations of 4-5% and no neuronal medium, with a second local maxima for 100% neuronal medium with 8-9% serum concentration. A region of low total cell fusion was observed for media containing neuronal medium with serum concentrations below 4%. The higher total fusion at serum concentrations above this region suggests that increasing serum concentration may compensate for the reduction in muscle cell quantity caused by the presence of neuronal medium.

**Fig 6.**
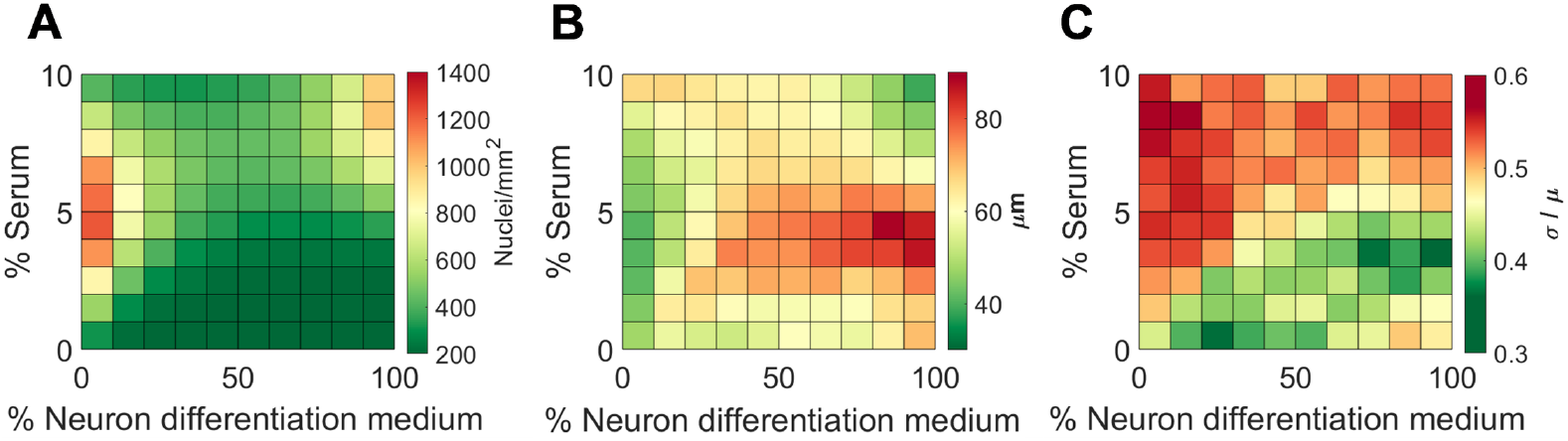
Surface plots showing predicted quality indicators over a range of media conditions from Agent Based Model output. (A) Difference in number of myonuclei between day 0 and day 5, (B) Mean distance between myonuclei at day 5 and (C) coefficient of variation of myonuclei at day 5.

Increasing the proportion of neuronal medium was shown to increase the mean distance between myonuclei (Fig 6B) with a maximum at a media composition of 4-5% serum and 80-90% neuronal medium. Predictions of the coefficient of variation (Fig 6C), show a decrease in the uniformity of myonuclei distribution as the concentration of serum increases and thus a predicted decrease in cell quality. Myonuclei uniformity did not decrease with increasing neuronal medium, indeed there are regions in which uniformity, and thus cell quality, is increased. The lowest variation in myonuclei distribution was shown to be in 100% Neuronal medium with 3-4% serum concentration with a further local minima at 20-30% neuronal medium and no serum.

### ABM simulations predict cell quality in 5% serum

To assess the predictive power of our model, we compared model outputs from our ABM against measurements of quality indicators from a further *in vitro* trial in media with 5% serum and 100% muscle cell differentiation medium. This media composition was chosen as our model(Fig 4) predicts local extremes of myonuclei density and coefficient of variation. We also compared these results with predictions from fitting a quadratic polynomial surface derived from averaged *in vitro* results. Paired t-tests indicate that there is no significant difference between ABM predictions and *in vitro* measurements (0.538 ±0.056 and 0.517± 0.25 respectively) for the coefficient of variation in myonuclei (Fig 7A), suggesting the simulations replicate uniformity of myonuclei distribution. ABM predictions outperformed a quadratic surface fit of *in vitro* results (Fig 7A), which predicted a coefficient of variation of 0.622, 20.3% higher than the experimental median.

**Fig 7.**
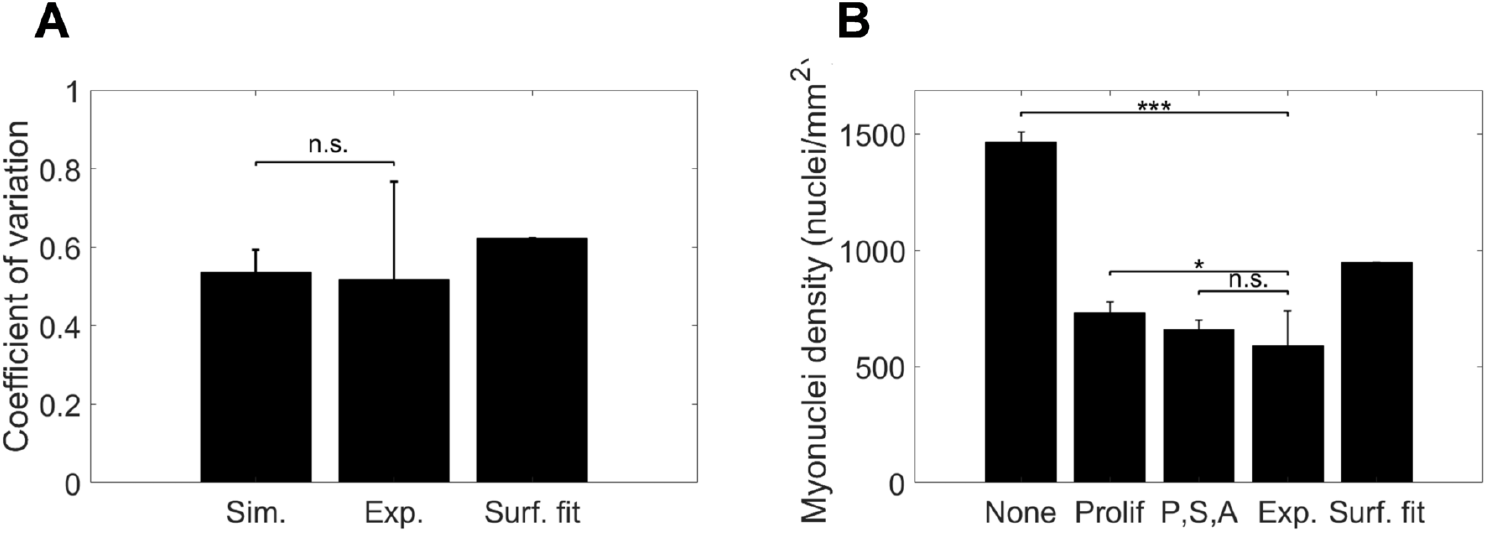
Comparison of simulated outputs of quality indicators against values measured from *in vitro* images. (A) Coefficient of variation (a measure of uniformity defined as the ratio of standard deviation to mean of distance between neighbouring myonuclei) in myonuclei comparing computational simulation against experimental measurement and prediction from a second-order surface fitted to averaged *in vitro* measurements. (B) Difference in myonuclei density between day 0 and day 5 for (from left to right); simulations with all behavioural inputs fitted from model, simulations with measured proliferation rate, simulations with measured proliferation rate, myoblast speed and myoblast angular velocity, experimental observation and prediction from a surface fitted model.

A significant difference was observed (Fig 7B) between ABM predictions of the difference in myonuclei density and *in vitro* observations (1466.8±40.5 and 589.2±150.8 respectively) To ascertain potential sources of model discrepancies, live images of cells in 5% serum with cell differentiation medium were analysed to observe cell behaviours and compare with those derived from surface fitting. Measured angular velocity and speed of cell motion exhibited modest deviations between fitted and measured (*mean absolute error* of 3.6 degrees/minute and 0.13 *μ*m/minute respectively), while cell proliferation rate was significantly lower than predicted (*Normalised mean absolute error* of 110%). Due to this large difference between predicted and measured proliferation rate, we exchanged the measured cell proliferation rate rather than the fitted value as an input to the ABM. This significantly reduced the difference between simulated and observed differences in myonuclei (Fig 7B). The additional application of measured values of speed and angular velocity further decreased the difference but with a much smaller effect. The myonuclei density predictions of the ABM with measured cell proliferation rate outperformed a quadratic surface fit of *in vitro* results (Fig 7B), which predicted 948 myonuclei per mm^2^, 61% higher than the experimental median.

### Computational modelling for all conditions can be completed in less than half the time required for a single *in vitro* trial

To evaluate the time saved using our workflow compared to non-computational methods of optimisation we break down the typical time taken to perform each step of the workflow. These timings are summarised in table 2

**Table 2.** Time taken to complete key workflow tasks.

| Task | Breakdown of timings | Total time | In parallel / additional to *in vitro* trials |
| --- | --- | --- | --- |
| Acquisition of live images | 2hrs/video for 7 trials. | 14 hrs | In parallel |
| Analysis of live images | 5 mins/video for 6 positions and 30 mins processing data for 7 trials. | 7 hrs | In parallel |
| Cell division count | 10 mins/video for 6 positions in 7 trials. | 7 hrs | In parallel |
| ABM-ABC computation of inferred behaviours | 20 mins/simulation for up to 300 simulations run in parallel on 16 cores for 7 trials. | 43.75 hrs | Additional |
| ABM Parameter sweep. | 20 mins/simulation for 121 simulations run in parallel on 16 cores. | 2.5 hrs | Additional |
| Total time | - | 74.3hrs | - |
| Total extra time | - | 46.3hrs | - |

Computational modelling in the workflow is designed for parallel computing and so the time taken to complete tasks will vary significantly depending upon the computational resources available. We assume that the *in vitro* trials required for alternative optimisation methods are at least as extensive as those proposed in our workflow.

Analysis of static, stained images took 2 hours for each trial (5 minutes to download, 5 minutes to pre-process images and 5 minutes to analyse per unique position for at least 6 positions with 30 minutes to collate information). Since analysis of static images is required for a typical *in vitro* analysis these timings have not been included here as an additional task for our proposed workflow.

Live images for each trial were acquired over a two-hour window. Automated analysis of myoblast motion behaviour for each trial took one hour (5 minutes of computation per unique position for at least 6 positions and 30 minutes to collate information). Manual counting of myoblast proliferation also took one hour (10 minutes per position). Live image acquisition and analysis can be run concurrently with *in vitro* trials.

A typical ABM run without the GUI takes between 10-20 minutes to simulate 5 days of cell differentiation and maturation. The ABC-SMC method requires up to 300 ABM trials to provide distributions of inferred cell behaviours. PyABC features provision for extensive parallel computing for this task and so any calculation of time required will depend upon the number processors available. We ran pyABC on a 16-core processor, taking a maximum of 6 hours 15 minutes to complete per trial (100 hours of processing time in total). Gaining inferred behaviour distributions for the 7 trials used here can therefore be achieved within a 2-day period.

Once calibrated, the ABM can then be used to predict cell quality and quantity for discrete media compositions (10-20 minutes per simulation) or sweep through the entire parameter space of media compositions. The parameter sweep conducted here of 121 discrete media compositions was achieved in under 3 hours using a 16-core processor to run simulations in parallel.

In total, the live image acquisition and analysis and computational effort required to implement this workflow was less than 3.5 days, of which up to 28 hours was run in parallel with *in vitro* cell maturation. For comparison, a single *in vitro* trial requires 9 days to reach day 5 of myotube differentiation (4 days in growth medium and 5 days in differentiation medium).

## Discussion

Optimisation of cell culturing protocols in tissue engineering is essential for maintaining the quantity and functionality of mature cells as well as ensuring experimental reproducibility. Applying quantitative mathematical and computational modelling approaches alongside *in vitro* experiments increases efficiency and reduces costs when designing cell culturing procedures, aiding the transition from bench to bedside for regenerative medicine [13].

This may take the form of a single early cell behaviour which correlates with quality indicators outcomes, allowing direct prediction, or may be a product of interactions between behaviours. Here, we apply an iterative *in vitro - in silico* workflow to culturing muscle cells from murine myoblasts with the aim of optimising concentrations of serum and neuron differentiation medium to maximise cell quality and quantity for a co-culture of muscle cells and motor neurons.

We employed metrics for quantifying the migratory and proliferative behaviour of myoblasts and developed novel metrics for indicating the quality and quantity of mature cells. We found that varying the concentrations of media components produces significant changes in indicators of mature muscle cell quality and key early-stage cell behaviours. We created and calibrated an agent-based model (ABM) to predict indicators of cell quality using metrics of early stage behaviour as inputs. Describing metrics of the cell behaviours as a function of media composition allowed the ABM to predict cell quality for any given concentration of serum or neuron differentiation medium without the need to measure further metrics of cell behaviours.

Combined *in vitro – in silico* experiments indicate that choice of media composition presents a balance between cell yield and cell quality. We show that increasing the concentration of serum in cell differentiation medium increases the total amount of myoblast fusion but has a detrimental effect on cell quality. We also show that cell quality can be maintained (or even improved) by the addition of neuronal differentiation medium though this also comes at the cost of lower myoblast fusion when compared to muscle cell differentiation medium alone. Uniform distribution of myonuclei is known to be associated with cells in healthy muscle tissue [14], with aggregation of nuclei linked to muscular disfunction [10] [15] [16]. In this study, the mean distance between myonuclei and coefficient of variation of myonuclei were applied as metrics of the uniformity of myonuclei distribution within a cell. The difference in myonuclei density per mm^2^ over the first 5 days of differentiation was used as a second indicator of cell outcomes.

Serum is added to media to stabilise conditions for cell proliferation though differentiation in some muscle cell lines has been shown to be triggered by the deprivation of serum. Serum represents an unknown in terms of composition, which may alter motor neuron differentiation efficiency and trigger loss of stemness. Our results show that an increase in media serum percentage leads to increasingly non-uniform distributions of myonuclei, a sign of less viable muscle cells. These findings agree with previous work by Lawson and Purslow [8] which shows C2C12 cells differentiated best in low serum or serum-free media and other studies [17] which show using media with low or no serum is advantageous in preparation for co-culturing muscle cells with iPS derived motor neurons. Our work, however, indicates that the relationship between serum concentration and cell fusion is non-linear, and that increasing serum to moderate levels (up to around 5%) increases the total amount of cell fusion. This could enable adding serum to muscle cells cultured in neuronal differentiation media to boost cell quantity without impacting quality. Trials showing greater increases in myonuclei density at the end of the differentiation phase exhibited a lower fusion index during the early stages of differentiation. This appears counter-intuitive, but since our simulations indicate that myoblast availability is a limiting factor in cell fusion rate, we hypothesize that media compositions with a lower fusion index prioritise myoblast proliferation over fusion during the early stages of differentiation. The supply of myoblasts will therefore not be exhausted as quickly as more fusion-efficient regimes, leading to a greater total amount of fusion. The negative relation between fusion index and total amount of fusion has wider implications for the reliance of fusion index alone as a marker of experimental success. As fusion index measures the ratio of myonuclei to mononucleated myoblasts at a given time, it conflates the rate of cell-cell fusion with the proliferation rate of myoblasts. While fusion index is simple to calculate and gives an intuition of initial experimental efficiency, it is important to consider that a high fusion index may be due to a high fusion rate of neighbouring cells or a low rate of background cell division, or some more complex balance between the two.

Since availability of myoblasts appears to be a limiting factor in the total amount of fusion, and thus the final volume of cells, it may seem intuitive to culture cells to confluence during the growth phase so there is greater availability. Previous studies [18] however, show that seeding myoblasts at a high density induces quiescence in cells and suppresses differentiation and so would not provide an effective strategy. Committed myoblasts need to switch from a proliferation state (growth phase) to a fusion state (differentiation phase). In *in vitro* culture, this switch is triggered by lowering media serum content which causes myoblasts to initiate cell cycle arrest and differentiate into myotubes, but this behaviour may be strongly altered if medium composition and serum percentage are not optimal. Calibration of our ABM highlighted a significant difference between measured cell proliferation rates and those predicted by a surface fitting model and from previous findings [8] that the cell proliferative state is preserved at higher serum concentrations.

Our numerical modelling assumes that metrics of cell motion do not change over time. It may be expected that cell migration speed and angular velocity change as myotube density increases with time though we found no significant changes in metrics of behaviour (results not shown) throughout the first day of differentiation. Temporal changes over a longer timescale were not studied but assumed to have a minor impact on model outputs due to the short timespan (1 – 3 days) in which the bulk of fusion occurs. Our model also assumes that all myoblasts may differentiate into myotubes. C2C12 cells which are deficient in the production of the proteins Myf-5 and Myo-D have been shown to fail to form myotubes in culture [19]. The low myonuclei densities observed here for high levels of neuron differentiation media indicate that not all myoblasts are viable. Nuclei stainings which differentiate between myoblasts and myonuclei would allow a clearer observation of myoblast fates under varying media conditions and allow for the quantification of non-fusing myoblasts into the model.

Additional early markers of cell quality include myotube width and the presence of actin striations in myotubes [20].We found no significant differences in width and a high proportion of actin striations present in all trials and so did not include them in our investigation though they may be relevant for optimising other cell culture protocols or cell lines.

Current practices in cell culturing tend to apply a trial-and-error approach to experimental design, though there is a call to move to more systematic methods of optimisation [5]. Non-computational methods for optimisation such as design of experiments (DoE) methods require interpolation of the results from *in vitro* trials. Our workflow does not seek to replace these methods, but to enhance their speed, cost and, potentially, accuracy as an iterative process between *in vitro* and *in silico* experiments.It also provides an insight into the mechanisms underlying the changes in cell quality which is absent in methods relying purely on regression analysis.

Our results suggest that predictions made by a calibrated ABM can outperform a quadratic surface model of *in vitro* results for measurement of spatial uniformity of myonuclei and for myonuclei density when the background proliferation rate of myoblasts is known. Myoblast proliferation rates can be extracted from live images during the first day of cell differentiation and so do not require cells to be cultured for long periods of time or for them to be stained. Despite this, the workflow would be more effective without a reliance on the measurement of proliferation rate to enable prediction. Further study of the effects of cell culture conditions on cell proliferation or adapting the environment to control rates of myoblast proliferation may provide a solution to the requirement for additional *in vitro* trials.

The workflow described here is designed to be applicable to other cell types, providing they exhibit early behaviours and quality indicators which can be quantified and that mature cell quality is not correlated with a single behavioural indicator. In this study we optimise just two cell culture conditions, but once behavioural and quality indicators have been acquired, adding further variables for optimisation is trivial. The number of trials required for optimisation using a DoE method increase exponentially with increasing numbers of factors observed and so the application of a workflow for reducing the number of *in vitro* trials will become increasingly more cost-effective. Factors for optimisation include further media compositions, materials for substrate structure and topological constraints such as patternings for aligning cells.

The design and calibration of the ABM from scratch is the most time consuming element of our workflow and requires specialist skills which may not be immediately available to labs. We have designed the ABM to be both user-friendly and customisable for general application, using open-source software packages where possible. NetLogo software for ABM design provides a user-friendly GUI. To obtain inferred cell behaviours, NetLogo is linked to PyABC via the PyNetLogo extension, example code available on Github, (https://github.com/dhardma2/MyoChip). PyABC features provision for extensive parallel computing for this task and so a calculation of the time required for calibration will depend upon the number processors available. By switching off the cell fusion mechanism, our ABM can be applied as a generic model of motile cell behaviour which could be used as a base for building similar predictive models of further cell types. While the fusion dynamics of the ABM could have potential in studying the formation of further types of syncytia, a more obvious application is the optimisation of further skeletal muscle cell lines including human induced pluripotent stem cells and the study of muscular degenerative diseases.

In summary, we identified new quality indicators for muscle cells, which can be implemented in future statistical approaches to optimisation. We showed that changes in the composition of muscle cell differentiation media affect both early-stage cell migratory behaviours, cell fusion index and cell quality. By representing early-stage cell behaviours as a function of media composition, we designed an ABM to predict cell quality for any given configuration of media. Our results indicate that the choice of culture media composition will involve a compromise between the quality of cells and total cell yield. The iterative workflow of *in vitro – in silico* experiments presented here can be applied as a tool for the optimisation of a wide range of further muscle cell culturing parameters and muscle progenitor cell lines, including human iPS, providing an efficient and cost-effective method for improving and quantifying tissue engineering procedures.

## Methods

### *In vitro* primary myotube culture

*In vitro* myotubes were cultured from primary mice myoblasts as described previously [21]. Shortly, hind limb muscles were isolated from 5-7 day old mice pups, minced and digested for 1.5 hours at 37°C using 0.5 mg/ml collagenase (Sigma) and 3.5 mg/ml dispase (Roche) in Phosphate-buffered saline (PBS). Subsequently, filtered cell suspension was plated in IMDM (Invitrogen) for 4 h in the incubator (37°C, 5% CO2). To purify cells from fibroblasts and other contaminating cell types, only non-adherent myoblasts were collected and centrifuged. Cells were resuspended in growth medium (IMDM + 20% FBS + 1% Chick Embryo Extract + 1% Penicillin/streptomycin) and plated onto 1:100 matrigel (RD) coated ibidi dishes. After 4 days, medium was changed to trigger differentiation. Depending on the experimental condition, standard muscle differentiation medium (IMDM + 1% Penicillin-Streptomycin) and neuronal medium (N2B27 medium: 50% DMEM-F12, 50% Neurobasal +1X N2 + 1X B27 + 50 *μ*M *β*-Mercaptoethanol + 1% Penicillin-Streptomycin) were mixed 1:0, 1:1 or 0:1, containing 0, 2, 5 or 10% horse serum. After 1 day of differentiation, a thick layer of 1:1 matrigel was added on top of forming myotubes and 100ng/ml agrin was added to the culture medium. Cells were cultured up to 7 days at 37°C/5% CO2.

The Rodent Facility of the Instituto de Medicina Molecular Joãao Lobo Antunes maintains high standards of animal welfare and promotes a responsible use of animals, hence supporting state-of-the-art animal based research. The Rodent Facility has a license of establishment for breeding and animal use, issued by the Portuguese competent authority - Direcçaão-Geral de Alimentaçaão e Veterinária (DGAV), since 2013. This facility complies with Portuguese law Decreto-Lei 113/2013, transposed from the European Directive 2010/63/EU, and follows the European Commission recommendations (2007/526/EC) on housing and care of animals and the FELASA (Federation of European Laboratory Animal Science Associations) guidelines concerning laboratory animal welfare. (Ethics approval number AEC 2014 03 EG)

### Live imaging

Live imaging was performed using an inverted microscope (Zeiss Cell Observer SD or Zeiss Cell observer) in widefield mode, using a 20x phase air objective (Plan-Apochromat Ph2 NA 0.80 or EC Plan-Neofluar Ph 2 NA 0.50, respectively). Cells were imaged 2 hours after adding the respective differentiation medium. 3×3 tiles with 10% overlap with 2×2 binning were taken every 5 minutes for a minimum of 12 hours. Images were stitched using Zen Blue software and stacked into 30 minutes videos for further quantitative analysis.

### Immunostaining and static imaging

Cells were washed once with PBS and fixed in 4% PFA for 10 minutes at room temperature. Cells were permeabilized (PBS + 0.5% Triton) for 5 minutes and blocked in blocking buffer (10% in Goat serum in PBS + 5% BSA) for 1 hour at room temperature. Primary antibodies (1:200; Anti-*α*-Actinin (Sarcomeric) mouse monoclonal antibody, Sigma Aldrich and A7732) were diluted in blocking buffer containing 0.1% saponine and cells were incubated at 4°C over night. Dishes were washed 2x for 5 minutes in PBS under agitation. Secondary antibodies (1:400; Goat anti-Mouse IgG Alexa Fluor 555, Thermo Fisher Scientific, A21424; Goat anti-Rabbit IgG Alexa Fluor 647, Thermo Fisher Scientific, A21245) in blocking buffer were incubated in presence of DAPI (100*μ*g/ml) for 1 hour. Dishes were washed twice, 200ul Fluoromount-G (Invitrogen) was added on top of cells. Samples were stored at 4°C. Dishes were imaged with an inverted fluorescent microscope (Zeiss Cell observer) using a 40x phase air objective (EC Plan-NeoFluar NA 0.75). z-stacks of 1 *μ*m were taken as 3×3 tiles with 10% overlap and 1×1 binning. Subsequently, imaged were stitched in Zen Blue.

### Agent-based model

NetLogo software (Wilensky, U. (1999). NetLogo.

(http://ccl.northwestern.edu/netlogo/. Center for Connected Learning and Computer-Based Modeling, Northwestern University. Evanston, IL.) was used to create a nuclei-centred agent-based model (ABM). Code is available on Github(https://github.com/dhardma2/MyoChip). A list of abbreviation used are included in Table 3). A pseudocode description outlining the rules of the ABM is illustrated in Fig S5).

**Table 3.**
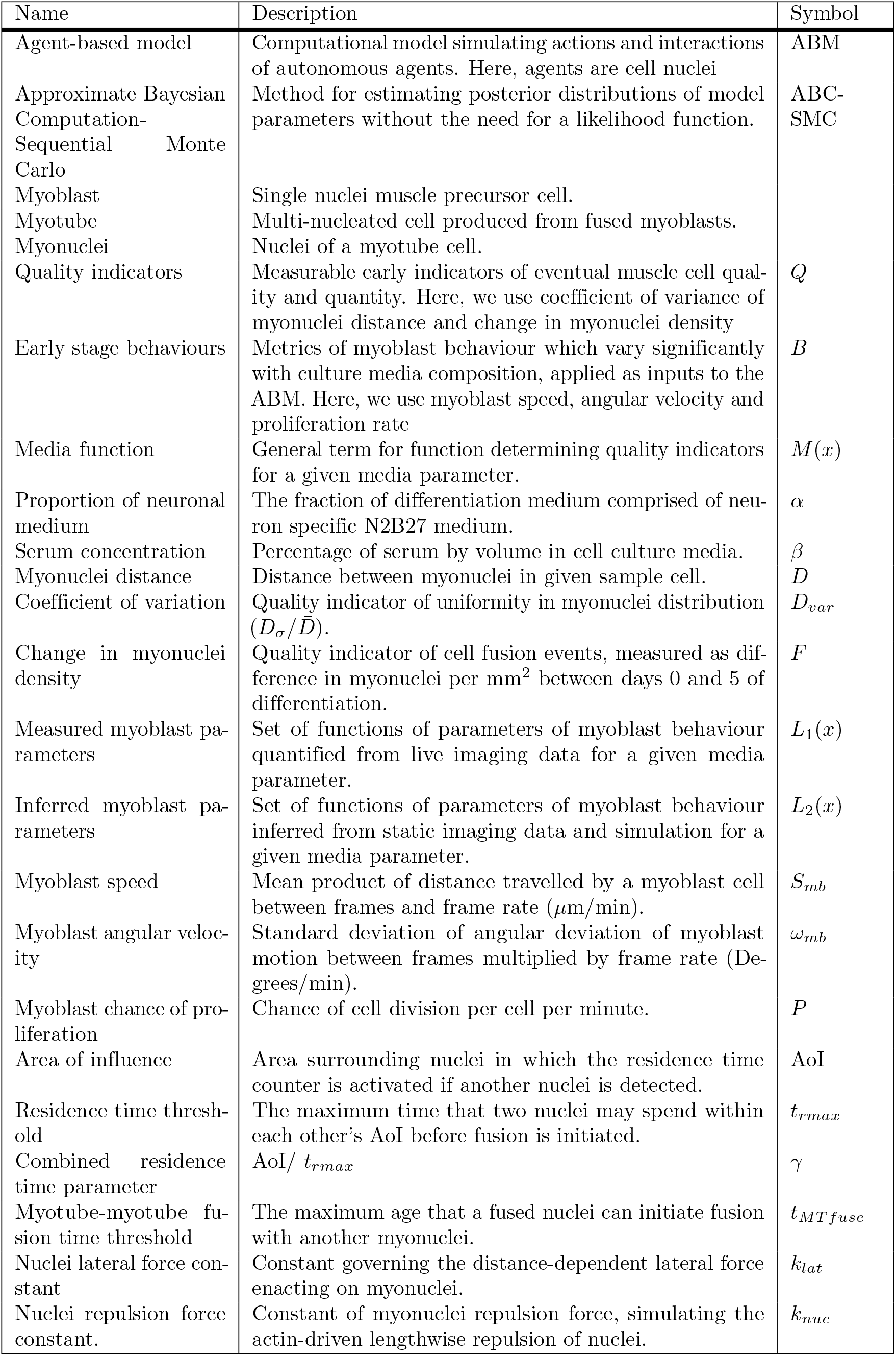
List of terms and abbreviations.

The ABM is seeded with initial densities of myoblast nuclei and myonuclei. Random sampling from normal distributions of measured and inferred myoblast behaviour metrics is used to apply the required inputs to model myoblast motion. Myoblast-myotube and myotube-myotube fusion is modelled via a residence time model. Once fused, a force balance is applied to myonuclei to simulate lateral and longitudinal motion within the cell. The ABM provides metrics of the densities of myoblast nuclei and myonuclei as well as distances between nuclei within discrete myotubes.

### Initialisation

The total number of seeded nuclei within the simulation is required for seeding, with nuclei initial position defined by random distribution within a simulated 1mm^2^ field of view. Each nucleus is initially assigned as a myoblast and given a probability of being converted to a myotube nucleus based on the inputted initial proportion of myonuclei. This element of stochasticity addresses the range of day 0 myonuclei densities recorded experimentally.

### Myoblast motion

At each timestep, myoblast nuclei are assigned a turning angle, chosen randomly from a normal distribution defined by inputs of mean and standard deviation of angular motion, and a distance in which to move in their new direction, also chosen from a normal distribution defined by mean and standard deviation of myoblast speed. Setting the mean turning angle to a non-zero value imparts a bias in myoblast direction of motion. Since no significant bias was found experimentally, the mean was set to zero. To model myoblast proliferation, at each timestep myoblast nuclei are assigned a probability of dividing calculated from an input for chance of division. Dividing myoblasts are stationary for a 20-minute period (a typical period for myoblast division observed *in vitro* after which a second nuclei is created with both exiting in opposite directions as was commonly observed *in vitro*).

### Fusion

Modelling of cell fusion events in the ABM is guided by the residence time of two or more cell nuclei of any type when in proximity (Fig 8A). A maximum residence time (*t*_*rmax*_) and surrounding Area of Influence (AoI) were prescribed to all nuclei. When the distance between two nuclei is less than the radius of the AoI, a residence time count (*t*_*r*_) is initiated for both nuclei. When *t*_*r*_ *> t*_*rmax*_ the nucleus changes its state from myoblast to myotube, indicating cell fusion. Sensitivity analysis of reasonable values of AoI and *t*_*rmax*_ revealed a strong linear relation between both parameters in predicting final myonuclei density (supplementary materials Fig S4). Variables *t*_*rmax*_ and AoI can therefore be condensed into a single variable, *γ* = *AoI/t*_*rmax*_ with units *μm*^2^ /minute. To simplify the choice of parameters, below we fix AoI to a circle with radius of 20 *μ*m (set as a value within the typical range of myoblast diameters measured from *in vitro* images) and vary *t*_*rmax*_. Myonuclei which are initially seeded or created via the fusion of myoblasts are labelled with a unique number to identify discrete cells. Subsequent nuclei fusing into these cells will share this identifying number.

**Fig 8.**
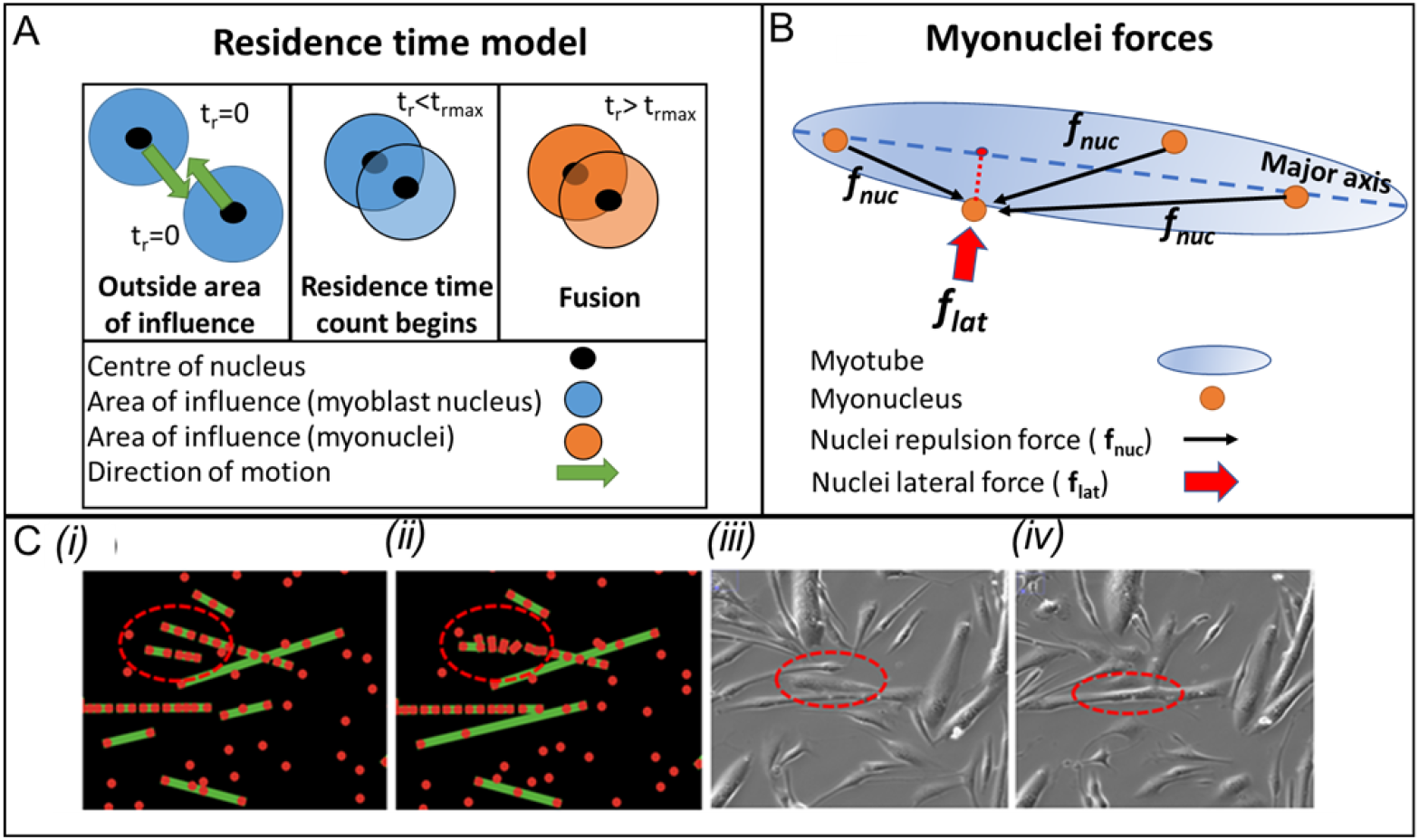
Overview of agent-based model. *(A) Residence time model of myoblast-myotube fusion. t*_*rmax*_ is threshold residence time. (B) Representation of forces acting on a single nucleus. A lateral force (*f*_*lat*_) moves the nucleus towards the major axis of nuclei in a myotube in a direction normal to the major axis. Total nuclei repulsion force is the sum of the individual repulsion forces (*f*_*nuc*_) acting on the nucleus. (C) Myotube-myotube fusion in agent-based model at (i) 0 minutes and (ii) 15 minutes (red dots represent nuclei and green lines represent myotubes) and in live images at (iii) 0 minutes and (f) 15 minutes.

### Myonuclei motion

To simulate the motion of fused myonuclei, a balance of forces acting on nuclei within each myotube was introduced to the ABM. Myonuclei have been shown to initially cluster at the centreline of the cell before spreading out laterally [10] [11]. Previous studies by Manhart *et al*. [22] [23] *on spatially constrained nuclei in Drosophila* myotubes successfully accounted for nuclei distribution over time by implementing a force balance. Forces considered in this study are (Fig 8b) a lateral force (*f*_*lat*_) to simulate the constraints myotube shape and a combined nuclei repulsion force *f*_*nuc*_ which simulates the actin-driven lengthwise movement of nuclei throughout the maturation process and any polar forces acting upon nuclei.

By denoting the position of the i-th nucleus at time t as the 2D variable X_*i*_(t), as characterised by Manhart *et al* [22], the velocity of the *i*-th nucleus is equal to the sum of all scalar forces acting upon it:

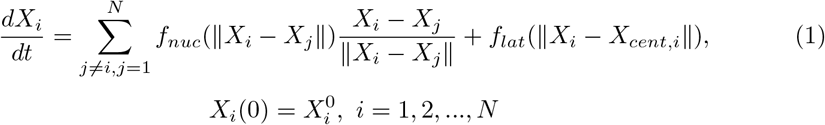

Where *X*_*cent,i*_ is the point on the major axis of nuclei in a given myotube closest to *X*_*i*_. The nuclei repulsion force can be expressed as:

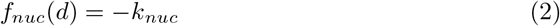

Where *k*_*nuc*_ is a value chosen from a normal distribution defined by a mean and standard deviation within the ABM and *d* is the distance between myonuclei. There is no limit on the distance at which nuclei can interact [23]. A similar force with magnitude inversely proportional to distance *d*, as described by Manhart *et al* was trialled, but was found to tend towards uniformity of nuclei distribution faster than described by our imaging data.

To apply *f*_*lat*_, a linear interpolation is applied to the co-ordinates of nuclei belonging to a given myotube to determine the major axis of the cell. The nuclei are then moved in a direction orthogonal to the major axis towards the centre of the cell with a magnitude:

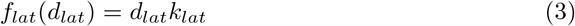

where *d*_*lat*_ is the shortest distance to the major axis from a given nucleus and *k*_*lat*_ is a constant defined within the ABM. During preliminary ABM simulations, nuclei of neighbouring myotubes were observed fusing together to create a single myotube (Fig 8c(i-ii)). This myotube-myotube fusion replicates observations *in vitro* (Fig 8c(ii-iv)) during differentiation. These fusion events continued throughout the entire span of differentiation in the ABM simulations resulting in myotubes containing significantly higher densities of nuclei than were observed *in vitro* at day 5 and beyond. From this we reason that there are mechanisms which constrain myotube-myotube fusion during the later stages of differentiation and so include a myotube-myotube fusion time threshold (*t*_*MT fuse*_) as a maximum ‘age-limit’ on fused nuclei after which they cannot fuse with other myonuclei.

### Outputs

At each timestep, the ABM records the number of myoblast nuclei and myonuclei per mm^2^, the fusion index (proportion of nuclei which are myonuclei), the mean and standard deviation of nuclei count per myotube and the mean 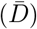 and standard deviation (*D*_*σ*_) of distances between myonuclei in each myotube. The coefficient of variation (*D*_*var*_), a measure of spatial uniformity of nuclei within a cell, is calculated as *D*_*σ*_*/*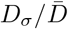).

## Model workflow

An overview of the workflow in creating a predictive model of cell quality is presented in Fig 9.

**Fig 9.**
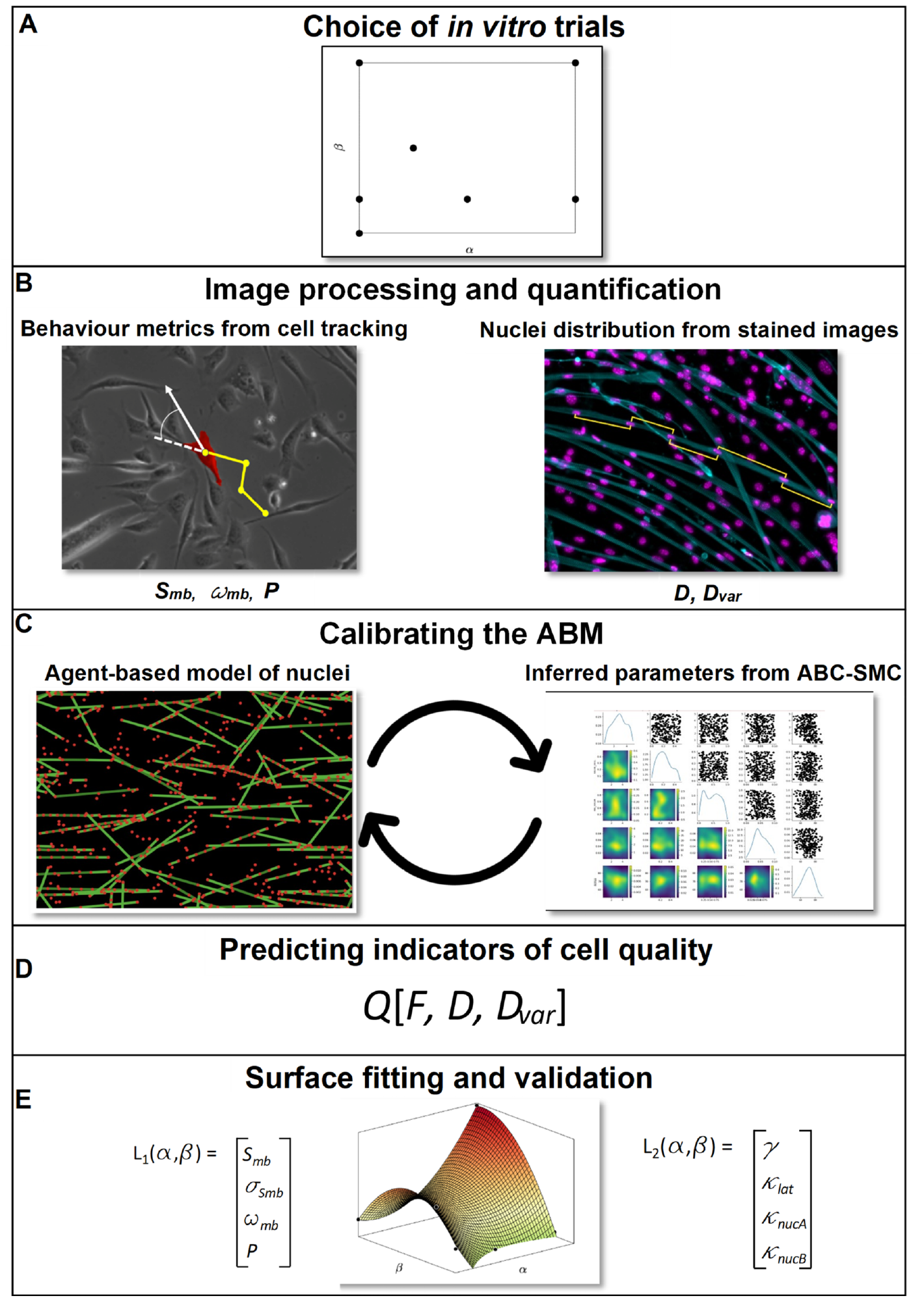
Overview of *in vitro-in silico* workflow. (A) Conduct *in vitro* trials with discrete values of media parameters *α* (Serum) and *β* (Neuron differentiation media).(B) Obtain measured distributions of myoblast speed (*S*_*mb*_), variance in angle of motion (*ω*_*mb*_) and proliferation rate (*P*) from tracking cells in live images. Calcuate density (*F*), mean distance (*D*) and coefficient of variation (*D*_*var*_ = *σ/D*) of myonuclei from fixed, stained images. (C) Representative images of ABM (left-hand side, red dots represent nuclei and green lines represent myotubes) and visualisation of posterior distributions from pyABC (right-hand side). Parameters of inferred cell behaviours derived using an Approximate Bayesian Computation method. Multiple ABM runs, informed by measured myoblast behaviours, are compared with measured quality indicators (*Q*). (D) Calibrated ABM predicts (*Q*) for a given set of early stage behavioural metrics (*B*) (E) To estimate ABM behaviour metrics for all values of *α* and *β* fit second-order surfaces to values of measured (function *L*_1_(*α, β*)) behaviours and inferred (function *L*_2_(*α, β*)) behaviours. Apply functions *L*_1_ and *L*_2_ to the ABM to estimate *F* and *D*_*var*_ for any value of *α* and *β*.Validate predictions against as further *in vitro* trial.

### Choice of *in vitro* trials

*In vitro* experiments with varying proportions of cell culture media variables were carried out to provide data with which to initialise, calibrate and validate the ABM (Fig 9A). We conducted seven trials (0% serum with 0:1 ratio of neuron differentiation medium, 2% serum with 0:1 ratio of neuron differentiation medium, 10% serum with 0:1 ratio of neuron differentiation medium, 2% serum with 1:1 ratio of neuron to muscle differentiation medium, 5% serum with 1:3 ratio of neuron to muscle differentiation medium, 2% serum with 1:0 ratio of neuron to muscle differentiation medium and 10% serum with 1:0 ratio of neuron to muscle differentiation medium) for validation of the model and a further trial (5% serum with 0:1 ratio of neuron differentiation medium) for model validation.

### Image processing and quantification

Live images of myoblasts are analysed to provide metrics of early-stage cell behaviours and fixed, stained images of myoblasts and myotubes provide cell quality indicators (Fig 9B).

Mononucleated myoblast cells were identified by cell area and segmented from live brightfield images (Fig 9 B) through a semi-automated process using an in-house MATLAB code. Cell centroids were tracked over two-hour periods providing data for calculating metrics of cell motion. Absolute values of the shortest distance between cells in consecutive frames were used to calculate myoblast speed. Angular velocity was calculated from clockwise or anti-clockwise change in angle of cell motion between frames. Applying Rayleigh-Moore tests to mean angular velocities showed that global cell motion showed no directional preference for all trials. The standard deviation of angular velocity, doubled to account for directionality, was applied as a metric of angular velocity over time.

Cell proliferation rate was determined by manual detection of cell divisions in ImageJ from live imaging over a two-hour period divided by the number of myoblast cells segmented in the first frame, giving a metric of chance of division per cell per minute.

The number of myotube and myoblast nuclei per mm^2^ were identified using MATLAB (R2019a) (MathWorks,Natick,MA) from static images with cell nuclei and actin cytoskeleton staining (Fig 9 B). Myonuclei distribution was measured in MATLAB by randomly selecting myotubes, manually labelling nuclei and calculating the distance between neighbouring nuclei. Coefficient of variation in myonuclei distribution can then be calculated from these measurements. Estimated mean myotube width and percentage of myotubes with actin striations were also recorded as potential indicators of cell quality, but not significant differences were found between measured experiments (See Supporting Information, Figures S1 and S2).

MATLAB code for segmentation and calculation of metrics is available on Github(https://github.com/dhardma2/MyoChip).

Behaviours and indicators which vary with changes in cell culture variables are retained. If no direct relationship between any single early-stage behaviour and cell quality metrics can be found then an ABM is employed to model the interactions between behaviours and ascertain whether predictions of cell quality arise as an emergent behaviour.

### Calibrating the ABM

Metrics of early-stage cell behaviours are applied as inputs for the ABM. Nuclei positions are the agents manipulated in the model. The main mechanisms of the ABM, chosen for their simplicity, are a residence-time cell fusion model and a force-balance model acting upon fused nuclei.

The ABM was calibrated by a comparison of simulated quality indicator metrics against those measured in discrete *in vitro* trials (Fig 9C). Metrics of cell and nuclei behaviours both observed and inferred (Table 4) from *in vitro* trials were applied as inputs to the ABM. Certain parameters required to drive these mechanisms could not be measured and so, where they have been identified, they are inferred by strategically comparing many ABM simulations with quality indicators using an approximate Bayesian computation sequential Monte Carlo (ABC-SMC) method [24]. We identified the parameters *γ* (nuclei area of influence / residence time threshold), *t*_*MT fuse*_, *k*_*lat*_ and *k*_*nuc*_ to be determined via statistical inference using the ABC-SMC method. ABC approximates a posterior distribution for each parameter by comparing the distance between summary statistics provided by the ABM with those measured from experimental images using a distance function (*δ*(*x*)). Parameter value *x* is rejected if the value of *δ*(*x*) exceeds a threshold distance (*ε*), building a posterior distribution from accepted parameter values. To speed this process, a sequential Monte-Carlo method was applied to systematically reduce the size of *ε* and so converge towards a choice of ABM parameters which best reproduce the experimental observations. The ABM-SMC method accounts for stochasticity within the ABM and provides information concerning uncertainties in parameter approximation. Inferred parameter distributions were calculated using the pyABC [25], an open source Python package well suited for parallelising tasks. NetLogo ABM simulations and pyABC were linked in Python using the pyNetLogo Python package. Examples of code applying the pyNetLogo interface are available https://github.com/dhardma2/MyoChip The number of fusion events (*F*), mean coefficient of variation (*D*_*var*_) and median myonuclei distance (*D*) at day 5 of differentiation were chosen as input summary statistics. *D*_*var*_ is a measure of the uniformity of myonuclei distribution but does not describe the magnitude of spatial distribution and so D is required to provide a sufficient measure of myonuclei distribution.

A distance function was defined as:

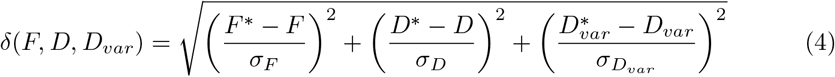

**Table 4.** ABM input parameters.

| Model parameter | Observed or inferred from data |
| --- | --- |
| Myoblast count at $t_0$ | Observed |
| Myotube proportion at $t_0$ | Observed |
| Myoblast speed of motion ($S_{mb}$) | Observed |
| Myoblast angular velocity ($\omega_{mb}$) | Observed |
| Myoblast proliferation chance ($P_m$) | Observed |
| Nuclei area of influence/Residence time threshold ($\gamma = AoI/t_{rmax}$) | Inferred |
| Coefficient of lateral force ($k_{lat}$) | Inferred |
| Coefficient of nuclei repulsion force ($k_{nuc}$) | Inferred |
| Myotube-myotube fusion time threshold ($t_{MTfuse}$) | Inferred |

This provides the *L*^2^ norm of differences between results from ABM simulations (*) and data measured from experimental images for *F, D* and *D*_*var*_, normalised by the standard deviations of measured data, giving dimensionless residuals for comparison. We assumed the posterior distribution for each parameter to be normal and found the mean and standard deviation as summary statistics for each experimental trial.

The posterior for parameter *k*_*lat*_ did not converge towards a fixed normal distribution. This is not surprising, as the arguments of the distance function are not direct indicators of the speed at which myonuclei move towards the centreline. Despite this lack of convergence, the posterior for all experiments suggested a range of feasible values for *k*_*lat*_ to be between 0.02 and 0.09. To represent this within the ABM, a value for *k*_*lat*_ between this range was randomly choosing from a uniform distribution at each required timestep.

If the ABM can be calibrated to simulate individual *in vitro* experiments then we assess its viability as a predictive model (Fig 9D).

### Surface fitting of early stage behaviour metrics

To determine the cell behaviour inputs for the ABM for any combination of differentiation media compositions without the need for further *in vitro* measurements, surface fitting was applied to the observed and inferred cell behaviour metrics obtained from discrete *in vitro* trials Fig 9E). A second-degree polynomial surface model was applied as suggested for efficient response surface methodology approximation by Box and Wilson [26]. Function subset *L*_1_(*α, β*), where *α* is the concentration of neuronal medium and *β* the percentage of horse serum in the culture media, is composed of functions of observable myoblast behaviour (myoblast speed, standard deviation of speed, standard deviation in angular velocity and chance of cell proliferation per minute) and function subset *L*_2_(*α, β*) is composed of functions of cell behaviours not directly observable but necessary to inform an ABM of myoblast-myotube cell fusion and dispersion (nuclei area of influence, residence time threshold, coefficients of nuclei lateral and repulsion forces and myotube-myotube fusion time threshold). The two subsets of functions were combined to create a composite function M(*L*_1_(*α, β*), *L*_2_(*α, β*)). A parameter sweep was then conducted through values from 0 to 100% neuronal medium and from 0 to 10% serum concentration using the function M(*L*_1_(*α, β*), *L*_2_(*α, β*), with early indicators of muscle cell quality *Q*[*F, D, D*_*var*_] as an output. Outputs were obtained from averaging over 10 simulations for each parameter set to account for stochasticity within the model. To ensure that all ABM simulations in the parameter sweep begin with similar cell seeding densities we use the mean of the experimentally measured nuclei densities of myoblasts and myonuclei day 0 of maturation to provide the seeding density and proportion of myonuclei, recorded as 420 and 53% respectively.

### Model validation

To assess whether the model predictions are accurate or whether measurement of some, or all, early-stage behaviours are required in order to make accurate predictions of cell quality a further *in vitro* trial was conducted for validation with 5% serum concentration and a 0:1 ratio of neuron to muscle differentiation medium. Quality indicator metrics were measured from static, stained, images and compared with those obtained via the ABM parameter sweep for the same media composition. Metrics of early stage behaviours were measured from live images taken during days 0-1 of this trial to compare predictions from the results of the parameter sweep with predictions made by including one or more measured early stage behavioural metric as an input. Second-degree polynomial surfaces were fitted to averages of the values of *F* and *D*_*var*_ measured from the *in vitro* trials to provide an example of interpolating cell quality results without the use of computational modelling as a baseline with which to compare the results of our workflow. Surface fitting was achieved using the *fit* function in MATLAB, using the fit type *poly22*. Second order fitting was used to account for non-linear relations but not overfitting the limited data points available.

### Statistical analysis

The Wilcoxon signed-rank test was applied to assess mean distance between myonuclei and two-sample t-tests were applied for all other data sets below unless otherwise stated. Statistical analysis was conducted in MATLAB.

## Supporting information

Supplemental FIgure 5, Pseudocode of ABM

Tupplemental Table 1, comparison of simulated and measured results

## Supporting information captions

**Fig S1. Comparison of change in average estimated myotube width over time in cells cultured in standard and neuronal differentiation medium** Width estimated by dividing total area of myotube cells by total length of myotube centrelines. No significant differences apparent between width of cells in different conditions.

**Fig S2. Comparison of proportion of myotube cells with actin striations in cells cultured in standard and neuronal differentiation medium** Data extracted from manual count from random sampling of cells stained for actin cytoskeleton. No significant differences apparent between proportion of striated cells in different conditions.

**Fig S3. Comparison of ABM and *in vitro* measurements of change in myonuclei density over time** for 2% serum and no neuronal medium.

**Fig S4. Relation between area of influence and residence time threshold** Blue circles denote local minima in a parameter sweep comparing difference in day 5 myonuclei density between ABM results and *in vitro* measurements for (a) 2% serum and no neuronal medium and (b) 2% serum and 100% neuronal medium. Red line represents least squares regression line.

**Fig S5. Pseudocode for agent-based model**

**Table S1. Comparison of calibrated ABM results and *in vitro* measurements**

## Acknowledgments

This project has received funding from the European Union’s Horizon 2020 research and innovation programme under grant agreement no. 801423. This research was funded in whole, or in part, by the Engineering and Physical Sciences Research Council (EPSRC) (grant nos. EP/R029598/1 and EP/T008806/1).

## Notes

https://github.com/dhardma2/MyoChip

### Competing Interest Statement

The authors have declared no competing interest.

### Summary of Updates

Revised to include detailed information on computational methods and improve figure quality as well as minor additions to the scope of the study.

