## Supplemental Figures S1-S4 for "An *in vitro* - agent-based modelling approach to optimisation of culture medium for generating muscle cells"

### Supplementary materials: Figures 1

March 29, 2022

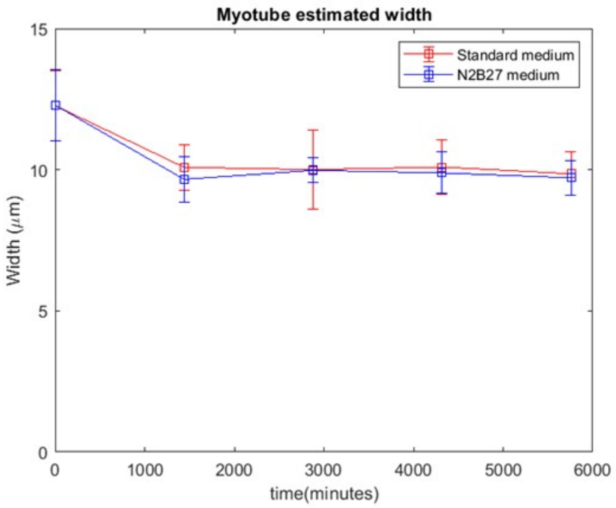


**Figure S 1. Comparison of change in average estimated myotube width over time in cells cultured in standard and neuronal differentiation medium** Width estimated by dividing total area of myotube cells by total length of myotube centrelines. No significant differences apparent between width of cells in different conditions.

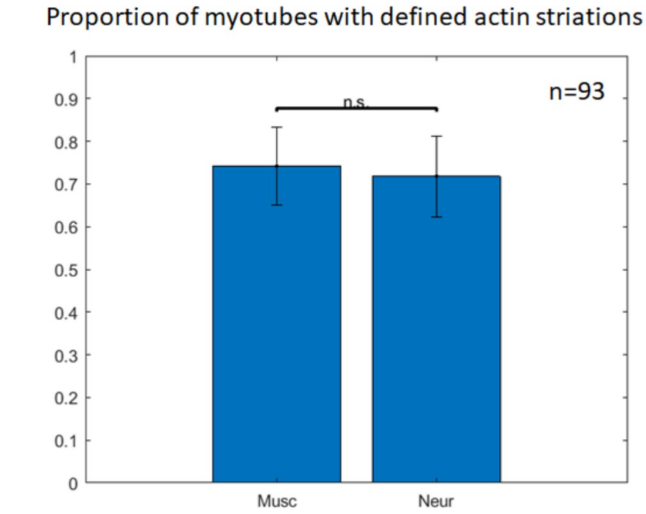


**Figure S 2. Comparison of proportion of myotube cells with actin striations in cells cultured in standard and neuronal differentiation medium** Data extracted from manual count from random sampling of cells stained for actin cytoskeleton. No significant differences apparent between proportion of striated cells in different conditions.

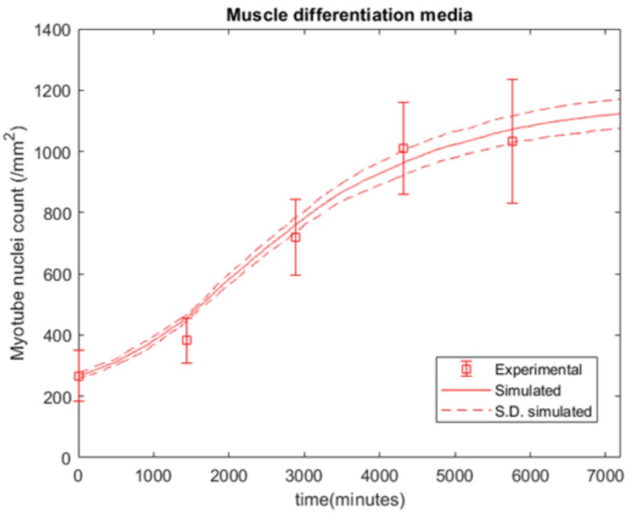


**Figure S 3. Comparison of Agent-Based Model and *in vitro* measurements of change in myotube nuclei density over time** For 2% serum and no neuronal medium

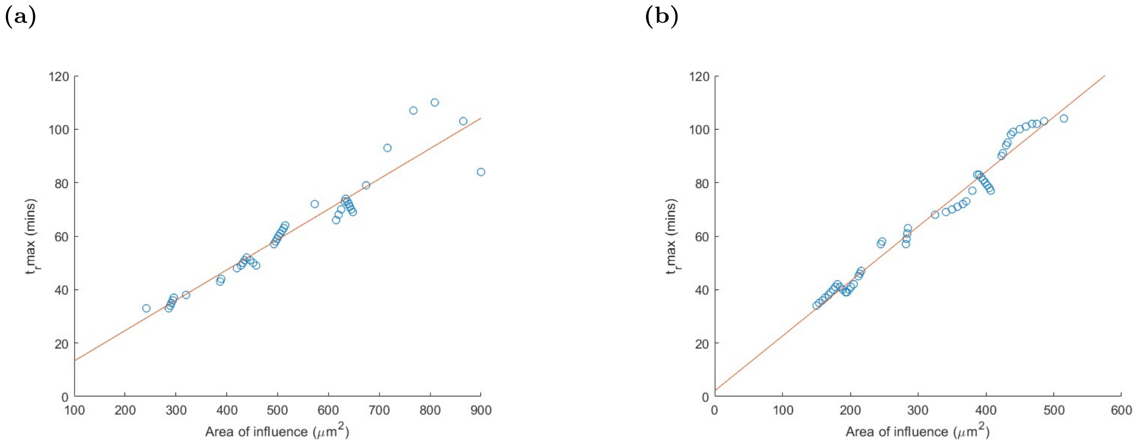


**Figure S 4. Relation between area of influence and residence time threshold** Blue circles denote local minima in a parameter sweep comparing difference in day 5 myotube nuclei density between ABM results and *in vitro* measurements for (a) 2% serum and no neuronal medium and (b) 2% serum and 100% neuronal medium. Red line represents least squares regression line.
