## Supplemental FIgure 5, Pseudocode of ABM for "An *in vitro* - agent-based modelling approach to optimisation of culture medium for generating muscle cells"

### Supplementary Materials : Figures 2

### Figure S5 Pseudocode for agent-based model

##### Initialisation

Seeding of *n* nuclei agents at random *x* and *y* coordinates

All agents assigned as myoblasts.

For each myoblast agent {

If randomly assigned percentage < input of myonuclei proportion

{Assign agent as myonuclei

Assign cell ID number *‘I’* }

}

##### Myoblast motion

While (simulation tick *t* < *T*) {

For each myoblast agent {

If randomly assigned number v [0,1] is less than (1/ *P*)

{Set agent proliferating = 1}

If agent proliferating = 0

{Choose angle from normal distribution with mean = 2*w_mb_

Rotate agent by chosen angle

Choose speed from normal distribution with mean = *S_mb_*

Move agent forward}

`	If agent proliferating = 1

{Increase proliferation tick + 1

If proliferation tick > 20

{Set agent proliferating = 0

Create new agent at same coordinates

Set angle of motion of new agent at 180^o^ to old agent}

}

}

}

##### Cell fusion

While (simulation tick *t* < *T*) {

For all agents{

If distance |agent_a_ - agent_b_| < radius *AoI*

{Increase residence time tick + 1 for agent_a_ and agent_b_

If residence time ticker > *t_rmax_*

{Assign agent as myonuclei

Set residence time ticker = 0

Agents share cell identifier *I* if assigned or assign new *I*}

}

}

}

##### Myonuclei dynamics

While (simulation tick *t* < *T*) {

For all myonuclei agents with cell identifier *I*{

Identify coordinates of agent

Apply linear regression to define function for centreline *C_I_* of cell *I*}

While myonuclei agents have cell identifier *I*{

For each agent{

Calculate *x* and *y* components of *k_nuc_* forces acting on agent

Calculate *x* and *y* components of *k_lat_* force directed towards *C_I_*}

Calculate magnitude and direction resultant vector of forces

Move myonuclei agent accordingly}

}

##### Data collection

If simulation tick *t* = *T*{

For all *I*{

Count number of myonuclei agents in cell *I*

For all agents in *I{*

Create single link between agent and neighbour in *I*

Measure distances between neighbours

}

Calculate mean and SD distance between myonuclei agents

Calculate coefficient of variation in myonuclei agents}}}}
