## Supplementary material for "An *in vitro* - agent-based modelling approach to optimisation of culture medium for generating muscle cells": Tupplemental Table 1, comparison of simulated and measured results

### Supplementary materials: Tables

**S1 Table 1. Comparison of calibrated ABM results and *in vitro* measurements**

| Differentiation media composition | | Difference in myotube nuclei density, day 5 (nuclei/mm^2^) | | | Coefficient of variation in myonuclei within a cell | | | Mean distance between myonuclei within a cell | | |
| --- | --- | --- | --- | --- | --- | --- | --- | --- | --- | --- |
| % Serum | % N2B27 | Experiment | Simulated | Difference from expt. | Experiment | Simulated | Difference from expt. | Experiment | Simulated | Difference from expt. |
| 0 | 0 | 345.3$\pm$57.4 | 338.6$\pm$21.8 | -1.9 % | 0.638$\pm$0.294 | 0.502$\pm$0.016 | -21.3% | 37.5$\pm$37 | 52.9$\pm4.7$ | +41 % |
| 2 | 0 | 846$\pm$274 | 767$\pm$34.4 | -9.3% | 0.583$\pm$0.267 | 0.541$\pm$0.047 | -7.2% | $47.7\pm32.7$ | $48.2\pm4.8$ | +1 % |
| 10 | 0 | 250$\pm$118.7 | 266$\pm$16.1 | +6.4% | 0.551$\pm$0.250 | 0.495$\pm$0.067 | -10.2% | $63.3\pm52.4$ | $60.1\pm5.6$ | -5.1 % |
| 2 | 50 | 19.6$\pm$55.4 | 268$\pm$3.8 | +1267% * | 0.450$\pm$0.241 | 0.446$\pm$0.045 | -0.9% | 60.4$\pm50.5$ | 66.4$\pm6.4$ | +10% |
| 2 | 100 | 56.4$\pm$98.1 | 272.8$\pm$4 | +384% * | 0.474$\pm$0.278 | 0.417$\pm$0.040 | -12% | 66.9$\pm34.2$ | 74.8$\pm3.8$ | +11.8% |
| 5 | 25 | 535.9$\pm$152.6 | 514.8$\pm$53.1 | -3.9% | 0.589$\pm$0.357 | 0.536$\pm$0.041 | -9.9% | 52.11$\pm42.45$ | 64.0$\pm6.0$ | +23% |
| 10 | 100 | 1464.3$\pm$265 | 1425$\pm$109.3 | -2.7% | 0.552$\pm$0.206 | 0.506$\pm$0.066 | -8.3% | 30.8$\pm18.4$ | 31.0$\pm1.43$ | +0.6% |

* ABM cannot model cases in which final nuclei count is less than the original seeded number of nuclei.
